# scatterbar: an R package for visualizing proportional data across spatially resolved coordinates

**DOI:** 10.1101/2024.08.14.606810

**Authors:** Dee Velazquez, Jean Fan

**Affiliations:** Center for Computational Biology, Whiting School of Engineering, Johns Hopkins University, Baltimore, MD 21211, USA; Department of Biomedical Engineering, Johns Hopkins University, Baltimore, MD 21218, USA

**Keywords:** data visualization, spatial transcriptomics, spatial analysis

## Abstract

**Summary:** Displaying proportional data across many spatially resolved coordinates is a challenging but important data visualization task, particularly for spatially resolved transcriptomics data. Scatter pie plots are one type of commonly used data visualization for such data but present perceptual challenges that may lead to difficulties in interpretation. Increasing the visual saliency of such data visualizations can help viewers more accurately identify proportional trends and compare proportional differences across spatial locations. We developed scatterbar, an open-source R package that extends ggplot2, to visualize proportional data across many spatially resolved coordinates using scatter stacked bar plots. We apply scatterbar to visualize deconvolved cell-type proportions from a spatial transcriptomics dataset of the adult mouse brain to demonstrate how scatter stacked bar plots can enhance the distinguishability of proportional distributions compared to scatter pie plots.

**Availability and implementation:** Scatterbar is available at https://github.com/JEFworks-Lab/scatterbar with additional documentation and tutorials at https://jef.works/scatterbar/.

## 1 Introduction

Effective data visualization plays a pivotal role in uncovering insights and communicating complex information clearly^1^. This is particularly important when dealing with proportional data in addition to spatial coordinates, as it combines quantitative values with their spatial context, providing a more comprehensive understanding of the underlying phenomena present and noting certain areas of interest. Such visualization of proportional data in addition to spatial coordinates is important for spatial transcriptomics (ST). ST enables high-throughput profiling of gene expression within tissue sections, often at multi-cellular pixel resolution^2,3,4^. Such data demand appropriate visualizations to accurately represent the presence of multiple cell types and their respective proportions in each pixel while preserving their spatial coordinates. Detecting cell-type-specific spatial variation such as proportional changes across spatial locations is important for understanding tissue organization and disease mechanisms. Therefore, there is a need for more effective visualization techniques that enhance the saliency of proportional changes and make it easier to interpret such complex spatial datasets.

Currently, a widely used method for this visualization is the scatterplot of pie charts, commonly referred to as ‘scatter pie plots’ as implemented through the scatterpie R package^5^. Scatter pie plots offer a way of displaying proportions at specific spatial locations by combining the spatial plane of scatterplots with the proportionate data representation of pie charts. However, in the context of visualizing ST data, scatter pie plots can leave significant amounts of whitespace in between spatial locations, which limits the efficient use of space, and are prone to visual artifacts such as moiré effects, which can distort perception. In general, the limitations of using pie charts to accurately visualize quantitative proportional information have been previously well characterized^6,7,8^. Specifically, pie charts encode quantitative data using angles, resulting in decreased accuracy in data interpretation compared to if the quantitative data was encoded using lengths^7,8^. Previous studies have shown that accurately comparing the sizes of the slices in a pie chart, especially when the differences are subtle, is more challenging than comparing the lengths of bars, for example, in a bar chart^7,8^. As such, the use of scatter pie plots for ST data visualization may make it challenging to accurately interpret and compare cell-type proportions to identify potentially interesting changes across spatial locations.

## 2 Materials and methods

Here, we present an R package called scatterbar, which provides an alternative visualization for displaying proportional data across many spatially resolved coordinates. Given a set of (x,y) coordinates and matrix of associated proportional data, scatterbar creates a stacked bar chart, where bars are stacked based on the proportions of different categories centered at each (x, y) location.

The package’s core functionality is in the function ‘scatterbar’. The function inputs comprise of a data frame (data) containing the proportions of different categories for each location, where each row represents a location in a 2-D plane and each column represents a category, and a data frame (xy) containing the positional information for each location specified in the row names.

To modulate the size of the bar plots, scaling factors (size_x and size_y) are used. Scaling factors are used to adjust the height and width of the bars relative to the plot’s coordinate system. They are crucial as they enable the visualization to fit within the specified plot area and thereby make the proportions more interpretable. If the scaling factors for the x and y axes are not provided, the optimal scaling factors are automatically estimated by computing the distance between the maximum and minimum values for each axis and dividing by the square root of the number of spatial spots. Padding for the x and y axes (padding_x, padding_y) can also be adjusted as needed. Padding refers to the additional space added around the bars to prevent overlap with the plot’s boundaries or other elements. Adjusting the padding ensures that the bars are clearly visible and separated from each other and the edges of the plot.

The final scatter bar plot is generated using the grammar of graphics framework as implemented in ggplot2^9^, with options for customizing the colors of the bars, the title of the plot, and the legend. The function returns a ggplot object representing the scatter bar plot, allowing users to apply standard ggplot2 functions like coord_flip() for orientation adjustments as well as ggplot2 extensions such as ggthemes for additional aesthetic modifications.

## 3 Usage scenario

To illustrate the potential utility of the scatterbar package, we created both the scatter pie and scatter bar plots to visualize a multi-cellular pixel resolution ST dataset of the adult mouse brain assayed by Visium^10.^ Briefly, for this dataset, gene expression measurements were profiled at 55µm resolution pixels across the tissue. Using these gene expression measurements, cell-type proportions per pixel were estimated using the reference-free deconvolution approach STdeconvolve^11^. The input to scatterpie and scatterbar thus contained the deconvolved cell-type proportions at each pixel and the corresponding spatial coordinates of each pixel centroid. As such, we aimed to visualize and compare the deconvolved cell-type proportions across spatial locations in the tissue.

While both scatter pie and scatter bar plots allow for the visualization of cell-type proportions across spatial locations, it can be more challenging to discern subtle differences in cell-type proportions between locations based on the scatterpie plot due to the visual similarity of angles in pie charts. In particular, we highlight two spatial locations where the proportion of green cell-type (i) is different, but the pink cell-type (j) is the same. In visually assessing the green and pink areas denoting the proportion of cell-types i and j respectively in the pie charts in the scatter pie plot, we note that it is difficult to discern whether these areas are the same or different across these two spatial locations in the tissue (Fig 1A). Alternatively, in visually assessing the green and pink bars denoting the proportion of i and j respectively in the bar charts in the scatter bar plot, we can more readily discern that the proportion of cell type i is different at the two locations but the proportion of cell type j is the same (Fig 1B).

**Figure 1.**
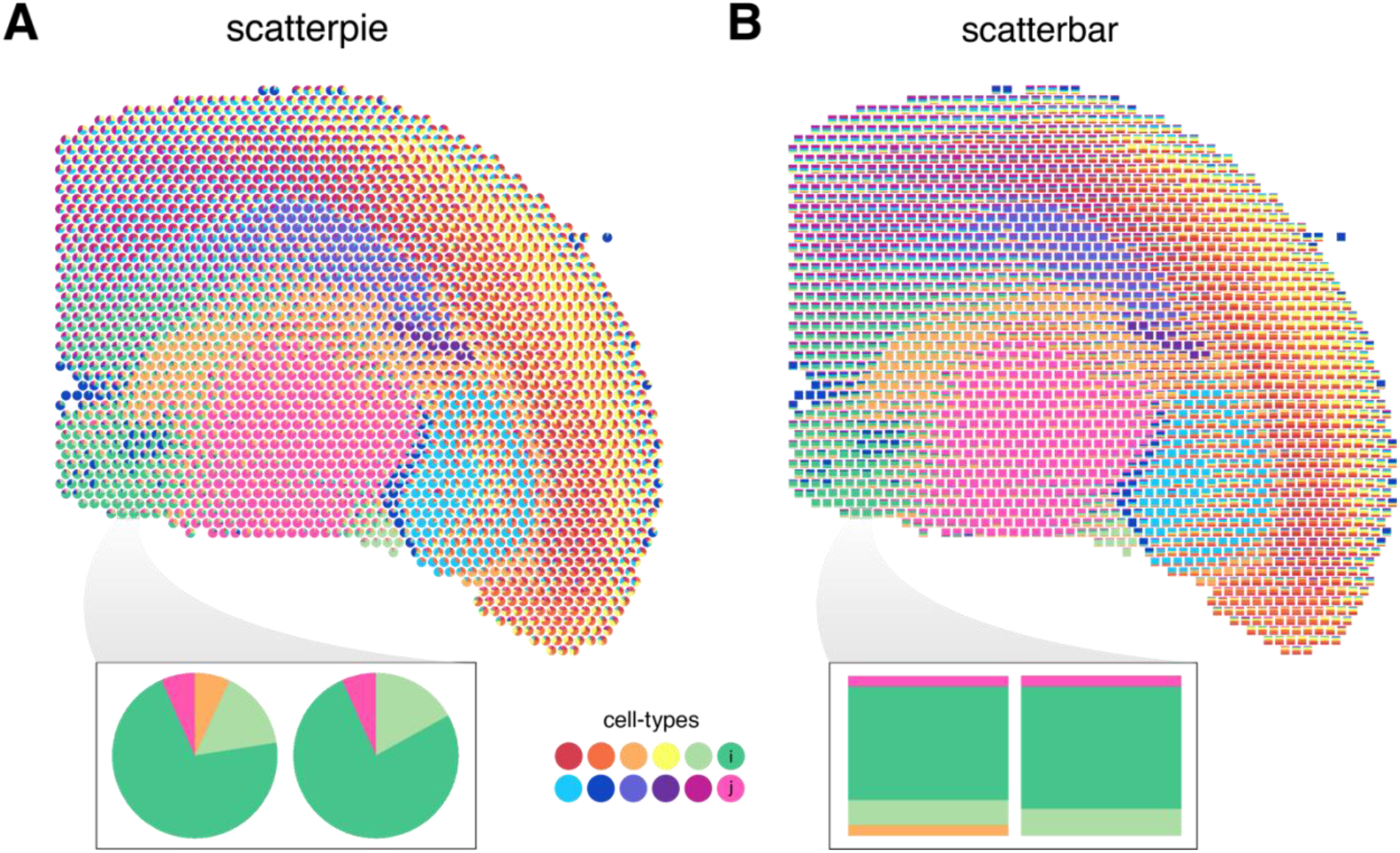
Comparison of scatter pie and scatter bar plots for deconvolved cell-type proportions in multicellular pixel-resolution spatial transcriptomics data of the adult mouse brain assayed by Visium. (A) Scatterpie plot showing the proportions of twelve cell types across spatial coordinates. Each pie chart represents the composition of cell types at that specific multicellular pixel in the tissue, with the size of the slice of each pie segment corresponding to the proportion of each cell type. (B) Scatterbar plot showing the same data. Each bar chart represents the composition of cell types at that specific multicellular pixel in the tissue, with the height of each bar segment corresponding to the proportion of each cell type. The x-axis and y-axis represent the spatial coordinates of pixels, and the colors correspond to the different cell types. Inset highlights two spatial locations.

By using bar charts instead of pie charts, scatterbar enhances the saliency of cell-type proportion differences between spots, making it comparatively easier to detect subtle variations in cell-type distribution across spatial locations in the tissue. Our observations align with the insights reported by researchers who have performed perceptual studies on data visualizations, reinforcing the effectiveness of the scatterbar approach in visualizing spatial transcriptomics data^1,6^.

## 4 Conclusion

We presented scatterbar as an R package implementing scatter bar plots for visualizing proportional data across spatially resolved coordinates. Compared to more commonly used scatter pie plots, scatter bar plots can enhance the saliency and distinguishability of proportional distributions, making it potentially easier to interpret and compare data across different, particularly neighboring spatial locations. By extending ggplot2, scatterbar allows users to readily customize and tailor the visualization to specific datasets and research needs.

Despite these noted enhancements and benefits, the effectiveness of scatterbar may still be limited by certain factors. First, scatterbar automatically calculates a scale factor to adjust the size of each stacked bar plot based on the data input. However, suboptimal visualizations may result when this scale factor is too large or too small. For example, when the scale factors for both axes are set too large, the stacked bar plots may overlap one over the other, making it impossible to discern all individual groups across the entire figure. Conversely, when the scale factor for either axes is too small, the stacked bar plots may become compressed, losing visual coherence and the ability to identify the groups and their proportions in an individual stacked bar graph. The padding between stacked bar plots also plays a significant role in both the readability of the plot and the perception of the data distribution. Excessive padding results in scattered and disjointed bars that obscure patterns as they overlap on top of each other (or should they be so excessive, result in no figure), reducing legibility. This demonstrates that appropriate padding depends on the scale and density of the data; autoscaling may not always provide satisfactory results. In general, we recommend users should experiment with padding to avoid under-or over-spacing and manually tune these various parameters to optimize their visualization.

Another critical limitation lies in color selection. Scatterbar relies heavily on distinct colors to differentiate between groups. In cases where user-selected colors are too similar, groups may become visually difficult to distinguish. As data complexity grows, and especially with larger numbers of groups, it becomes essential to carefully choose contrasting colors to maintain visual clarity. We recommend that users manually input their own color palettes if the auto-generated rainbow color palette based on the number of groups present in the dataset results in indistinguishable colors. We note that this challenge is not unique to scatterbar and approaches previously developed for color palette optimization may be integrated^12,13^.

Finally, by default, scatterbar orders the groups as they appear in the dataset. However, manual reordering may improve readability. Generally, in a stacked bar chart, the top and bottom bars are aligned with a consistent baseline, making them easier to visually compare across locations. In contrast, middle bars lack a common baseline, and their positions shift depending on the size of the bars below them, rendering visual comparisons more challenging due to the lack of a common reference point. In such scenarios, reordering groups by reordering the input data frame column ordering such as the first and last columns correspond to the groups of interest may facilitate comparisons. Such reordering could also facilitate interpretation in cases where the sequence of groups follows a meaningful hierarchy. For example, groups with typically higher proportions across the dataset could be ordered first to facilitate easier comparison across spatial locations.

In general, as the number of groups and the number of spatial locations increase, the task of directly comparing numerous proportions over extensive spatial scales becomes progressively more challenging. Alternative summary statistics and visualizations beyond those enabled by scatterbar may still be needed to address such challenges.

Although we have focused on demonstrating the utility of scatterbar in the visualization of deconvolved cell-types in a ST Visium dataset, it is applicable to other multi-cellular pixel resolution ST technologies such as Slide-seq^3^ and DBiT-seq^4^ as well. Likewise, although we have focused on demonstrating applications to ST data, accurately visualizing changes in proportional data is crucial for a variety of applications, including but not limited to geographic information systems, phylogeography, in addition to biological research. As such, we anticipate that scatterbar will be a relevant tool for a wide range of applications.

## Acknowledgements

This material is based upon work supported by the National Science Foundation under Grant No. 2047611. We thank Mayling Chen, Caleb Hallinan, Srujan Singh, and Kamil Slowikowski for testing and feedback of the software package.

